# Involvement of 5’ and 3’ UTRs on SARS-CoV-2 Genome Packaging

**DOI:** 10.1101/2024.09.19.611578

**Authors:** Zhang Zhang, Kun Yang, Fangze Shao, Wenlong Shen, Ping Li, Yue Zhang, Junjie Xu, Guoying Yu, Jun Zhang, Zhihu Zhao, Yan Zhang

**Author notes:** To whom correspondence should be addressed: Yan Zhang Correspondence may also be addressed to Zhihu Zhao,., Jun Zhang or Guoying Yu. The authors wish to note that the first two authors contributed equally. Competing interests: The authors declare no competing interests.

## Abstract

The SARS-CoV-2 virus, responsible for the COVID-19 pandemic, packages its large single-stranded RNA genome through a precise yet enigmatic mechanism. A packaging signal (PS) within its genome was proposed to facilitate the assembly of new viral particles. Here in this study, we report the role of the 5’ and 3’ untranslated regions (UTRs) in PS-mediated genome packaging. Utilizing proximity ligation data, we demonstrate direct interactions between the UTRs and the PS9 element, a key packaging signal within the SARS-CoV-2 genome. Multiple evidence confirmed that the presence of UTRs enhance the packaging efficiency of infectious virus-like particles (iVLPs) and recruits more nucleocapsid (N) protein, suggesting a critical role in genome compaction and packaging. These insights into the regulatory mechanisms of SARS-CoV-2 genome packaging provide novel targets for antiviral therapeutics and contribute to the broader understanding of coronavirus assembly.

## Introduction

Coronavirus is a type of enveloped virus possessing remarkably large single-stranded RNA (ssRNA) genomes ranging from 27 to 32 kilobases in length. These viruses typically exhibit spherical or pleomorphic particles with a diameter of 60-140 nanometers, characterized by club-shaped projections from the viral surface[1–3]. A pivotal phase in the coronavirus life cycle is the initiation of genome packaging, a process that entails the compaction of the viral genome by structural proteins and its subsequent assembly into nascent virions. A central enigma in this process pertains to the mechanisms by which coronaviruses discern and condense their expansive RNA genomes into more compact forms[4].

Coronaviruses,like many other single-stranded RNA (ssRNA) viruses, assemble their nucleocapsid (N) protein around the genome to form a beads-on-string ribonucleoprotein (RNP) structure[5–7]. The viruses genomic RNA contains specific RNA elements known as packaging signal (PS), which interact with nucleocapsid protein[8, 9]. Interactions between the nucleocapsid (N) protein and the packaging signal (PS) within the viral genome are crucial for initiating the packaging process and for excluding non-genomic RNA from the assembly[10]. Although the precise location of the packaging signal within SARS-Cov-2 genome is a subject of ongoing debate, one prominent candidate has been identified within a region designated as “T20” (nt 20080-22222). This T20 could be further truncate to “PS9” region (nt 20080-21171), both could promote virus assembly independentlys[8].

Previous research has indicated that coronavirus packaging is facilitated by packaging signals that induce phase separation of the nucleocapsid (N) protein, prompting speculation on the mechanisms governing viral packaging[11, 12]. These studies have also proposed the potential involvement of untranslated regions (UTRs) in N protein phase separation, thereby suggesting a role for UTRs in the coronavirus packaging process. Nonetheless, direct functional evidence substantiating the participation of UTRs in this process remains elusive[13–15]. To investigate the potential involvement of genomic element in SARS-CoV-2 packaging and to elucidate the underlying mechanisms, we leveraged proximity ligation data to delineate the RNA interactome associated with PS9[16, 17]. This lead to identification of interactions between the 5’UTR and 3’UTR with PS9, which is corroborated by simplified SPLASH and COMRADES data analysis. Subsequently, we studied 5’UTR and 3’UTR effect in infectious virus like particles (iVLPs) model since its modular design recapitulated virus assembly process initicated by packaging signal.

Our research has revealed an enhancing role for the 5’UTR and 3’UTR in PS9-mediated packaging, suggesting that these regions may play a critical role in the assembly of the viral genome.In our investigation into the mechanisms of UTR elements in the initiation of viral packaging, we identified a core sequence within these elements through an analysis of the SPLASH data. This analysis suggests that the interaction between the UTRs and the packaging signal 9 (PS9) is pivotal to the initiation process. Additionally, we assessed the relative expression levels of the nucleocapsid (N) protein within infectious virus-like particles (iVLPs). The presence of untranslated region (UTR) elements within the iVLP genome substantially augments the recruitment of nucleocapsid (N) protein, thereby implying a mechanism by which UTRs may enhance the packaging process of SARS-CoV-2.

We anticipate these results contribute to the fundamental knowledge of coronavirus assembly, and also hold promise for the development of therapeutic interventions aimed at disrupting this critical viral process.

## Results

### SARS-Cov-2 packaging signal spatially co-locolized with 5’UTR and 3’UTR

The fragment PS9, locating in nsp15 and nsp16 in SARS-Cov-2 genome, was reported as the viral packaging signals[8]. Drawing inspiration from the work of Syed et al, a virus like particles genome was cloned to validate its function. It is shown that although the iVLP genome is much shorter than SARS-Cov-2, PS9 significantly initiate viral structure proteins assembly to form spherical particles in about 100nm diameter, which is similar to the authentic viruses. Morphology and immunity similarities between SARS-Cov-2 and its iVLPs confirmed PS9 function in virus packaging initiation (Supplementary figure 1) and the potential of iVLP to be basic model of study packaging process.

To uncover the potential regulatory mechanisms underlying SARS-CoV-2 packaging initiation, we employed PS9 as a bait in our analysis of proximity data derived from the published SPLASH[16] and COMRADEs {Ziv, 2020 #3026}dataset. This approach allowed us to map the cis-active elements on the viral genome. Our findings revealed that both the 5’UTR and 3’UTR exhibited a higher frequency of interactions with PS9, particularly during the early phase of infection when the viral genomes are predominantly in a relaxed conformation (Figure. 1A). As the infection progresses, the specificity of these interactions may be diminished due to the general compaction of the viral genome, which could mask the specific binding events that are so apparent in the early phase.A similar analysis utilizing the COMRADES dataset corroborated our observations, demonstrating an analogous interactome between the UTRs and PS9 (Figure. 1B). These data collectively suggest that both UTR regions specifically interact with PS9 and may play a role in modulating the packaging process.

**Figure 1.**
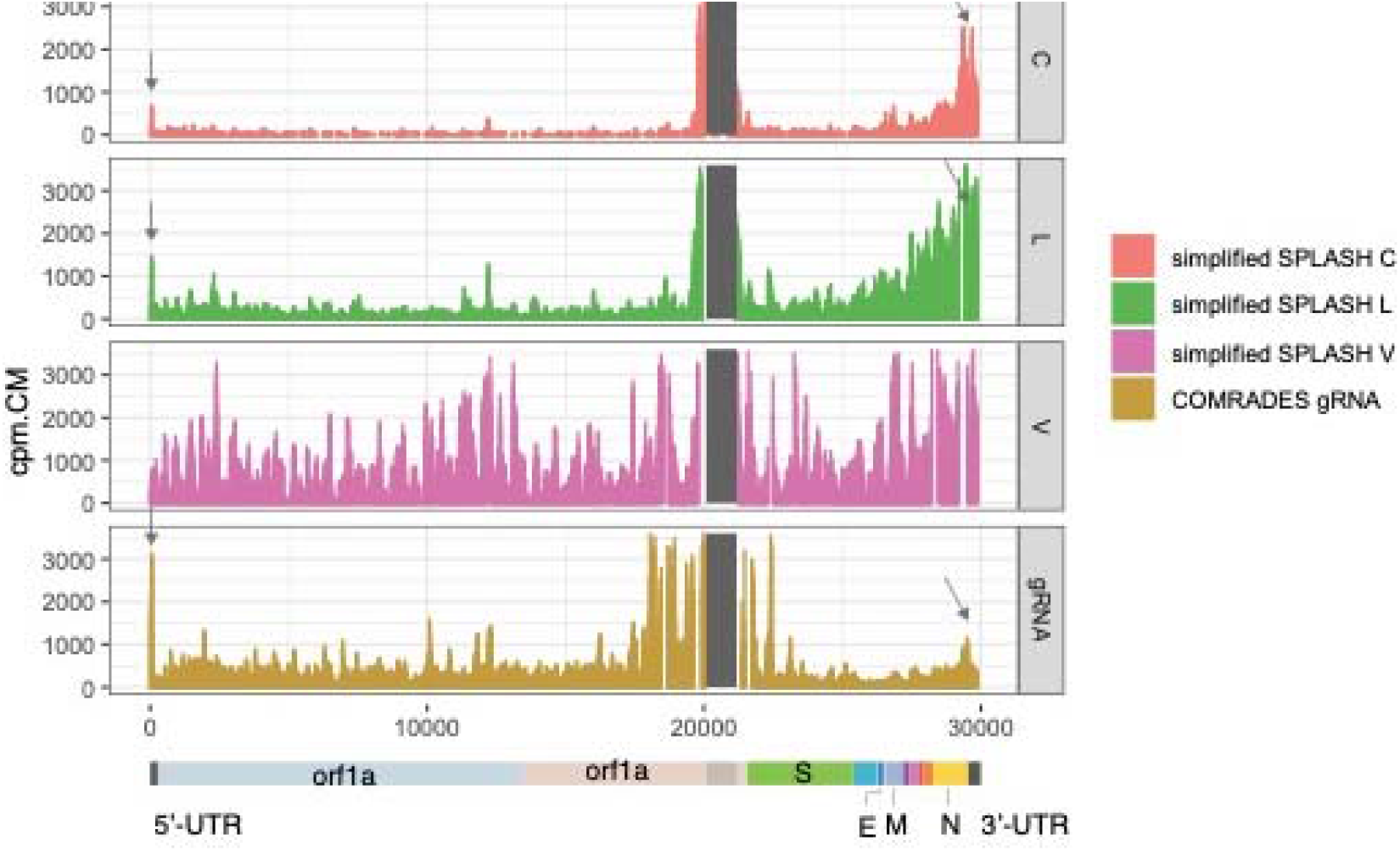
Proximity analysis of published SPLASH and COMRADES data. SPLASH data were captured in early infected stage (red histogram in C row), late stage (green histogram in L row), and virions (magenta histogram in V row). COMRADES data were captured in gRNA (ochre yellow). The shadow indicate the location of PS9 and the arrows indicate the interactive sequences. The bottom is diagram of SARS-Cov-2 genome to map the relative position of the sequences.

### 5’UTR and 3’UTR enhance the virus packaging efficiency

To delineate the role of UTRs in PS9-mediated SARS-CoV-2 packaging, we constructed a series of simplified, modular, and replication-incompetent iVLP genomes, with or without the incorporation of UTRs of SARS-CoV-2, to assess their impact on the packaging process (Figure.2A). These constructs were transfected alongside structural protein genes, and RNA from the supernatant of the resulting iVLPs was subjected to reverse transcription and droplet digital PCR (ddPCR) analysis.

**Figure 2.**
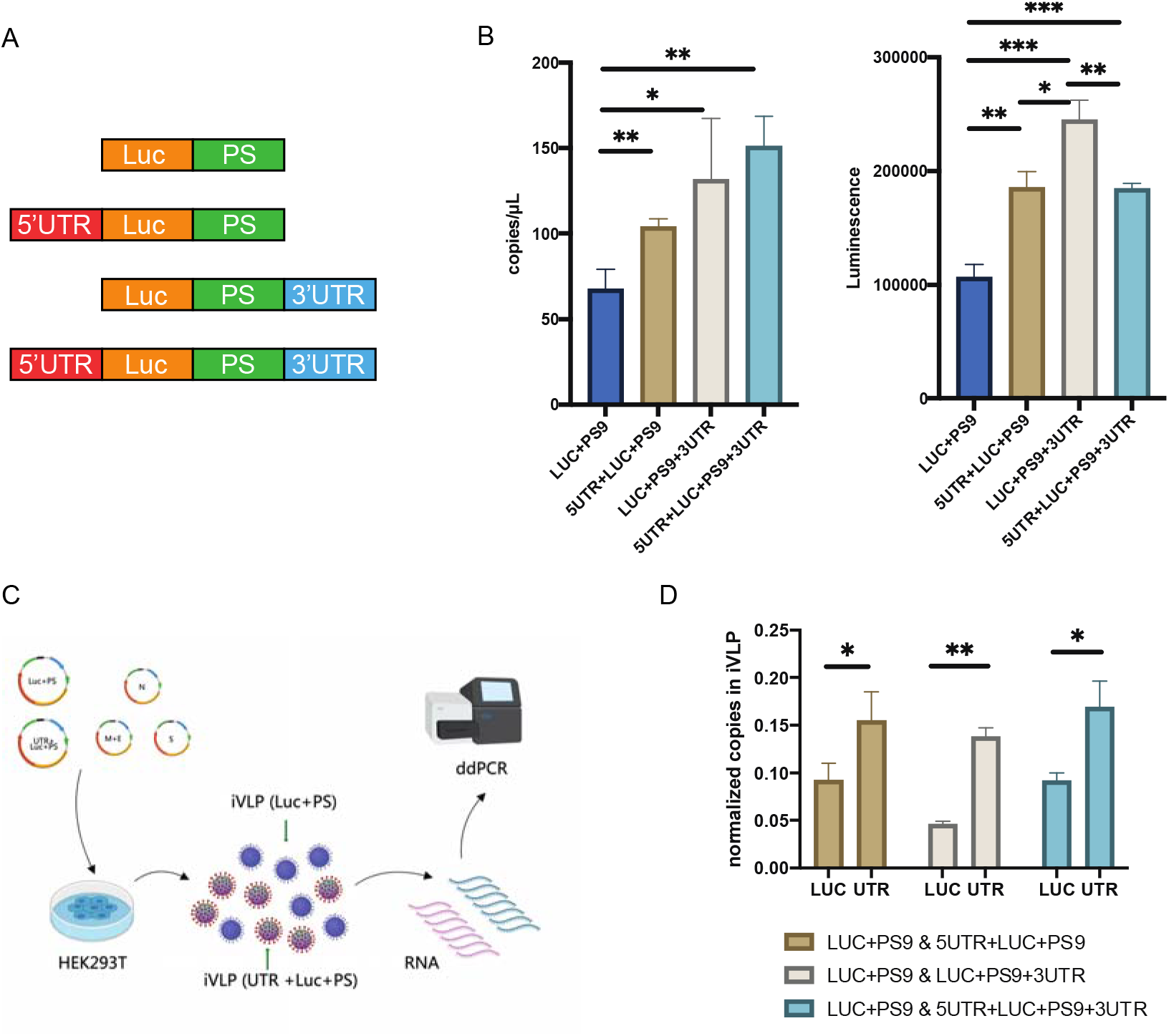
5’UTR and 3’UTR effect on SARS-Cov-2 packaging initiation. (A) Design of iVLP genomes. Luc, luciferase coding gene. PS, validated packaging signal PS9. 5’UTR and 3’UTR, 5’UTR and 3’UTR of SARS-Cov-2 genome. (B) Genome copies and luciferase readout of different genomes packaged iVLPs. (C) Overview of competition test of iVLP genomes with or without UTRs. (D) Normalized genome copies of different genomes in iVLPs.

The iVLPs containing SARS-CoV-2 5’UTR and/or 3’UTR exhibited higher copy numbers than iVLP lacking UTRs, indicating more viral particles in these samples.

Additionally, luciferase assays using ACE2-expressing HEK293 cells infected with RNaseA-treated iVLPs showed that the presence of UTRs significantly enhanced iVLP infectivity, aligning with the ddPCR findings (Figure 2B). Notably, iVLPs containing the 3’ UTR (Luc-PS-3’UTR) demonstrated the most pronounced effect. The infectivity of these iVLPs was further quantified using a nonlinear regression model to calculate the EC50 values from serial dilution infections, with Luc-PS-3’UTR iVLPs showing the lowest EC50, indicative of the highest infectivity (Supplementary Table 1). Additionally, luciferase readout from the ACE2-expressing HEK293 cells infected with RNaseA-treated iVLPs demonstrated that the presence of either the 5’UTR or the 3’UTR significantly enhances iVLPs infectivity, consistent with corresponding ddPCR results (Figure. 2B). Notably, And iVLPs containing 3’UTR (Luc-PS-3’UTR) displayed the most significant effect. The infectivity of these iVLPs was further quantified using a nonlinear regression model to calculate the EC50 values from serial dilution infections, with Luc-PS-3’UTR iVLPs showing the lowest EC50, indicative of the highest infectivity (Supplementary Table.1).

To further confirm the enhancing effect of UTRs on the packaging efficiency of virus particles, a pairwise competition assay was designed to directly compare functionality of UTRs. As illustrated in Figure. 2C, structural protein genes and two iVLP genomes were co-transfected in the same packaging system. The hypothesis is that a plasmid carrying the packaging signal, whether with or without UTRs, would exhibit a competitive advantage in packaging initiation, leading to a higher proportion of iVLPs in the final product. After confirming the amplification specificity of the primers and probes using qPCR (Supplementary figure.2), RNA from iVLPs supernants were extracted and subjected to ddPCR analysis. By normalizing transfection and PCR efficiency using trasfected plasmids across different primer sets, we demonstrate that iVLPs containing 5’UTR and/or 3’UTR produced a higher yield of iVLPs (Figure.2D). At the meantime, Luc-PS-3’UTR occupied the largest part in whole particles, which is consistent with iVLPs infectivity test results (Supplimentary figure 3). This finding indicates that the iVLP genomes with UTRs have a comparative advantage in the packaging process.

### 5’UTR and 3’UTR recruit more nucleocapsid proteins in iVLPs

It has been reported N protein could assembly in the presence of many viral RNA fragments, including PS9, 5’UTR and 3’UTR. It also has been reported that N protein could drive liquid-liquid phase separation, which could be promoted by 5’UTR and 3’UTR[11, 13, 14]. Given these considerations, we presumed that the integration of UTR elements into iVLP genomes might alter the N protein incorporation. To test this hypothesis, we detected the N protein content within iVLPs using Western blotting. S protein expression were used as internal control, since its not evolved in packaging signal interaction.

Interestingly, the results indicated that the content of the N protein was elevated in iVLPs containing UTRs. It is displayed that, compared to iVLP without UTRs, N protein content were elevated more than 2 folds in iVLPs with UTRs (Figure. 3A and Supplimentary figure. 4). To confirm this result, a digital western with replicants were performed as independent experiments, and the results showed that iVLP with UTRs were constitute of about 2 fold N proteins as well, which is consisitent to western blotting (Figure. 3B). The varied content of N protein expression suggested that UTRs recruit more nucleocapsid proteins in iVLPs assembly, which might be a direct result of 5’UTR and 3’UTR facilitate N assembly, or boosted liquid-liquid phase separation offer a convenient scope for virus assembly. It also supported that N protein is an critical trans activating element in SARS-Cov-2 packaging initiation.

**Figure 3.**
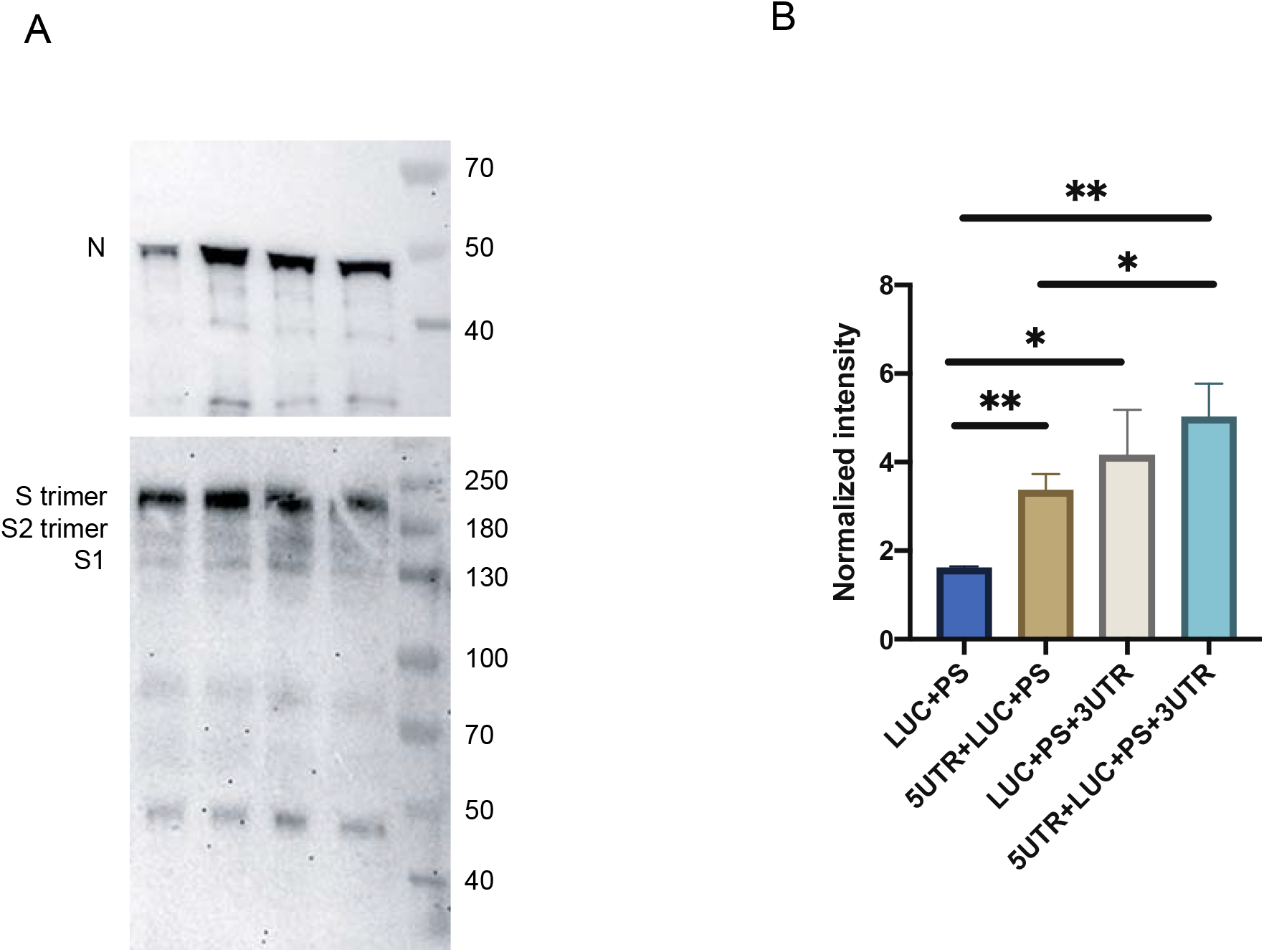
5’UTR and 3’UTR influence on relative N expression in iVLPs. (A) Western blotting detection the N and S protein in iVLPs. (B) Relative N expression detection in iVLPs by using digital western blotting.

### Identification of core sequence at 5’UTR and 3’UTR in virus packaging

We conducted an in-depth analysis of the SPLASH data. to elucidate the mechanism by which UTRs contribute to virus packaging, This analysis identified approximately 20 nucleotides within the 5’UTR and 30 nucleotides within 3’UTR that directly interact with PS9 (Figure 4A). We hypothesized that the deletion of nucleotides We hypothesized that the deletion of these interacting nucleotides might reduce the UTRs’ capacity to enhance packaging.. To test this, we created three truncated iVLP genomes: 5’UTRΔ49, which retains a putative interaction sequence despite the deletion of the first 49 nucleotides from the 5’ UTR; and 5’UTRΔ105 and 3’UTRΔ84, designed to eliminate PS9 binding sites (Figure. 4B). As anticipated, the iVLPs with deletions in the untranslated regions, specifically 5’UTRΔ105 and 3’UTRΔ84, demonstrated reduced luciferase expression levels. In contrast, the iVLP 5’UTRΔ49, which retains the critical interaction sequence, did not exhibit a significant reduction in luciferase expression relative to the iVLPs with complete 5’ and 3’ untranslated regions (Figure 4C). A pairwise competition assay further confirmed that 5’UTRΔ105 reduce its enhancement efficiency in packaging significantly (supplimentary Fig. 5), These findings underscore the importance of specific Watson-Crick base pairing between the UTRs and the packaging signal for the initiation of viral assembly, rather than the mere spatial proximity of UTR sequences to the packaging signal.”

**Figure 4.**
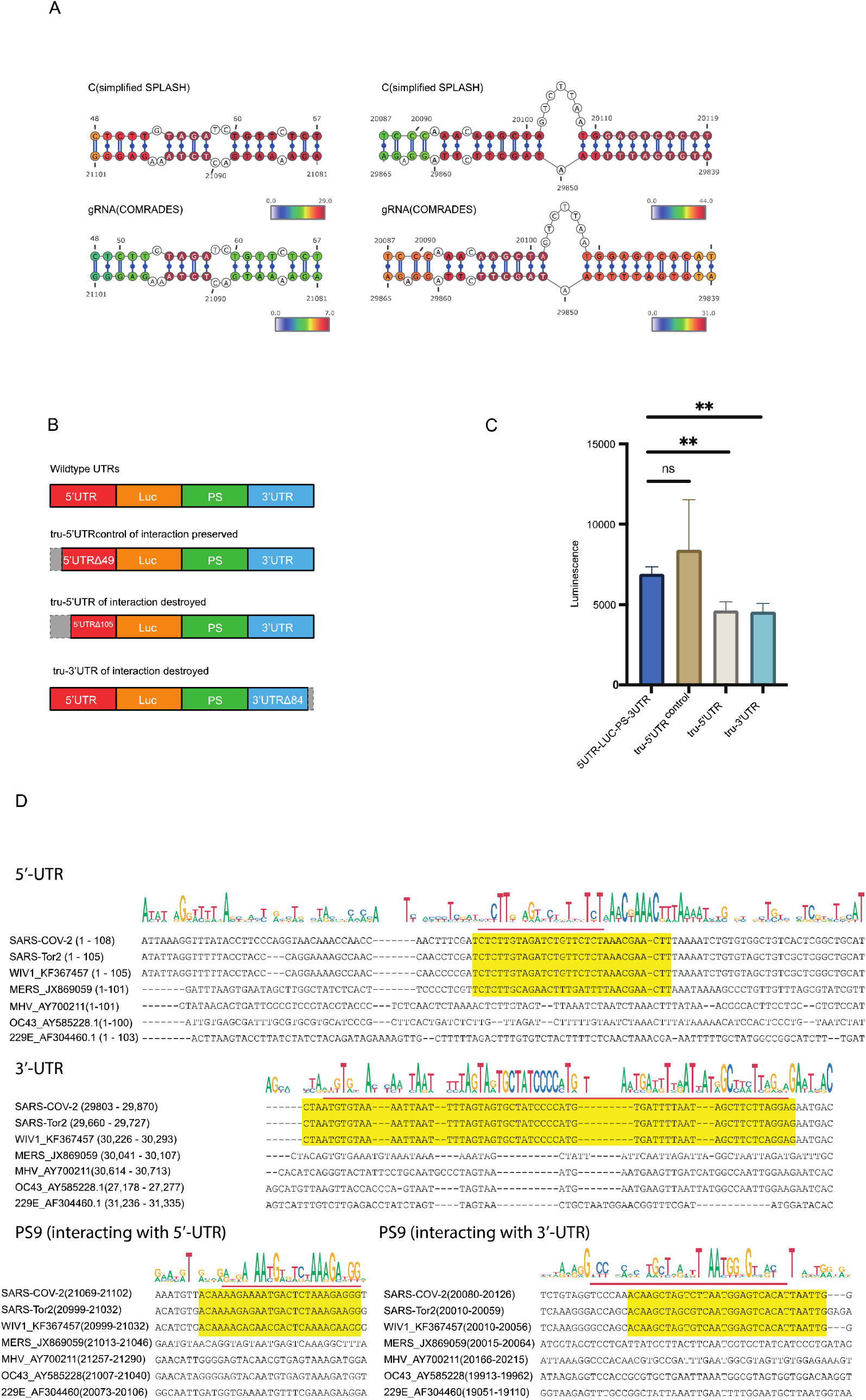
5’UTR and 3’UTR core sequence identification. (A) Base-paring analysis of UTRs and packaging signal in SPLASH and COMRADES data. (B) Design of iVLP genomes containing wildtype UTR or truncated UTRs. (C) Luciferase readout of different genomes packaged iVLPs. (D) Multiple sequence alignment of the core regions within the packaging signal (PS9), 5’ and 3’ untranslated regions (UTRs) of SARS-CoV-2 and several other coronaviruses.

Packaging signal, 5’UTR and 3’UTR core sequence within SARS-Cov-2 was multiple alignment with several famous coronavirus is performed. The results shown in figure 4D demonstrated high conservation in core sequence of these three regions, especially among SARS-Tor2 and WIV1, which have already coursed serious pandemic or poised for human emergence.(Fig.4D). We therefore proposed that interaction of UTR with PS9 might contribute to virulence of coronavirus. suggested that beside SARS-Cov-2, UTRs’ promoting effect on packaging initiation might be a prevalent mechanism in beta coronavirus.

We performed a multiple sequence alignment of the core regions within the 5’ and 3’ untranslated regions (UTRs) of SARS-CoV-2 and several other prominent coronaviruses. The results demonstrated significant conservation in the core sequences of both the 5’ and 3’ UTRs, especially among severe coronaviruses such as SARS-CoV-2, SARS-CoV, and SARS-like WIV1-CoV(WIV1), as well as Middle East respiratory syndrome coronavirus (MERS). These coronaviruses have either instigated substantial pandemics or are considered high-risk for human transmission[18]. In contrast, mild coronaviruses, exemplified by HCoV-OC43 and HCoV-229E, exhibited lower conservation levels. Our findings imply that the interaction between UTRs and the packaging signal 9 (PS9) could be a determinant of virulence in coronaviruse.

## Discussion

Our study unveils the intricate interaction between the SARS-CoV-2 packaging signal PS9 and its 5’ and 3’ UTRs, adding a sophisticated layer to our understanding of the viral packaging machinery. Evidence presented in this study substantiates the active involvement of UTRs in the packaging mechanism, as revealed by our analysis of proximity ligation data from the SPLASH and COMRADES datasets. This molecular insight suggests that UTRs directly engage with the PS9 element, participating in the assembly process.

Prior research has indicated that phase separation of the N protein, potentially triggered by packaging signals, is a key driver in viral packaging. Our findings support the hypothesis that UTRs participate in this phase separation, contributing to the ongoing scholarly debate on their role in coronavirus packaging.

Our results highlight the pivotal function of UTRs in augmenting the packaging efficacy of iVLPs by directly interacting with PS9 This enhancement appears to stem from a direct influence on the assembly of the nucleocapsid (N) protein, a critical component for the compaction and organization of the viral genome. Previous studies have postulated that the phase separation of the N protein, triggered by coronavirus packaging signals, might be a key driver of viral packaging[14, 18]. Our findings support the hypothesis that UTRs participate in this phase separation, contributing to the ongoing scholarly debate on their role in coronavirus packaging.

Despite the specific roles of the 5’ and 3’ UTRs in SARS-CoV-2, particularly in terms of gene translation and host-pathogen interactions, have been diversely interpreted[19–22], it’s suggested that UTRs lack the capacity to autonomously initiate viral particle packaging. This has led to an ongoing scholarly debate on the actual involvement of UTRs in coronavirus packaging. The present study advances the understanding of the contribution of UTRs to SARS-COV-2 packaging by demonstrating their role in facilitating the recruitment and assembly of the nucleocapsid (N) protein into viral particles, thereby elucidating a key aspect of the SARS-CoV-2 packaging mechanism.

Through truncation assay, we fine mapped essential regions that contribute to enhance packaging efficiency by interacting with PS9, which is consistent to base pair predition. This further surport the notion that UTRs contribute to cronovirus packaging by interacting with PS9. Interestingly, these core region is highly conserved among severe coronavirus but not in mild coronavirus. These prompt us to suspect that these interaction might confer virulence of cronovirus, which await further investigation.

It is important to note that previous study reported the mere act of truncation does not account for the observed decrease in assembly efficiency[8]. The differential capacity of RNAs, despite similar lengths, to induce N-protein condensation suggests that sequence and structure are pivotal in driving the assembly process[23]. Our study, therefore, provides a nuanced understanding of the multifaceted roles of UTRs in the packaging of SARS-CoV-2, offering insights that could inform the development of targeted antiviral strategies.

## Methods

### Cells

The Lenti-X 293T cell line was purchased from Clontech (now TAKARA Bio, USA), while the HEK-293T-ACE2 cell line was procured from Shanghai Anwei Biotechnology, CHINA. Cells were cultured in Dulbecco’s modified Eagle’s medium (DMEM, Pricella) supplemented with 10% fetal bovine serum (FBS, Pricella), 100 U/mL penicillin and 100 μg/mL streptomycin (Invitrogen, USA) at 37 °C in an atmosphere of 5% CO2.

### Plasmid Construction and Molecular Cloning

Plasmids Luc-PS9, M+E (CoV2-M-IRES-E), GFP-PS9, N (CoV2-N-R203M), and S (CoV2-Spike-D614G) were sourced from Addgene. ULP denotes the insertion of the 5’ untranslated region (UTR) of SARS-CoV-2 at the 5’ end of Luc in LP, while LPU represents the insertion of the 3’ UTR of SARS-CoV-2 at the 3’ end of PS9 in LP. ULPU signifies the insertion of both the 5’UTR of Luc in LP into the 5’ UTR of SARS-CoV-2 and the 3’ end of PS9 into the 3’ UTR of SARS-CoV-2. Variants such as 5UTRΔ49, 5UTRΔ105, and 3’UTRΔ84 involve targeted nucleotide deletions while preserving crucial interaction sequences.

### PS9 Interaction Identification

This study utilized various public datasets, including the simplified SPLASH dataset established by our laboratory, COMRADES data, and RIC-seq data. The original data was aligned to the SARS-CoV-2 reference genome. Utilizing the viewpoint method, the PS9 region was isolated, and interacting fragments across the entire genome were identified. This approach aimed to elucidate the comprehensive interactions of the PS9 region with the genome, laying the groundwork for further functional investigations. Additionally, based on the interaction’s intensity, determined by the number of supporting chimeric reads, long-range interaction bases were predicted. The mfold algorithm was employed to forecast RNA interactions, pinpointing specific paired bases.

### iVLP Production

Plasmids were co-transfected as manufacturer’s application note of improve lentiviral production using lipofectamine 3000 reagent (Invitrogen, USA). For 6-well culture plates, approximately 1.6 × 10^6 cells per well were seeded in 2 mL of lentiviral packaging medium. For 10 cm dishes, 9.6 × 10^6 cells per dish were plated in 12 mL of lentiviral packaging medium. Plasmids, lipofectamine 3000 and P3000 reagent were added according to instructions. After 48 hours, cell supernatant was harvested from each well or dish, clarified by centrifugation, and filtered using 45 µm pore size PES filter (PALL, USA). The clarified lentiviral supernatant was aliquoted into cryovials and stored at -80°C.

### Luciferase Assay

In each well of a transparent 96-well plate, 50 µL of supernatant containing iVLP was dispensed, followed by the addition of 100 µL of antibody-free 293T cell-specific medium containing 35,000 recipient cells (HEK-293T-ACE2). The plate was then incubated at 37°C with 5% CO2 for 24 hours to facilitate viral entry and infection. Following the incubation period, all the culture medium was carefully aspirated from each well of the transparent 96-well plate. Subsequently, 100 µL of Bio-Lite detection reagent (Vazyme), pre-equilibrated to room temperature, was added to each well. The plate was left at room temperature for a minimum of 3 minutes to allow for complete cell lysis. Thereafter, 80 µL of cell lysate from each well was transferred to an opaque white 96-well plate, and luciferase luminescence was quantified using a microplate reader Infinite M200PRO (Tecan, USA) with auto-attenuation function set and an integration time of 1000 ms.

### Competitive Packaging Experiments

During the packaging of iVLPs, besides the plasmids coding structure protein of SARS-Cov-2, the genome plasmid with or without UTRs was added in equivalent molar ratios. Supernatants were then collected for RNA extraction from the packaged virus-like particles, and the differential transcript content was assessed using droplet digital PCR. RNA extraction from virus-like particles or cells was executed utilizing Qiagen’s RNeasy Plus Kit. Quantification of specific gene transcripts was achieved through droplet digital PCR (ddPCR) using the Digital LightCycler 5× RNA Master Kit (Roche, swiss) with specific primers and probes. Amplification efficiency and specificity of primer sets were determined by qPCR of serial diluted template. Additionally, reporter plasmids were transfected and quantified simultaneously to normalized potential variation in transfection efficiency.

### Dual Luciferase Competition Experiment

During the packaging of iVLPs, besides the plasmids coding structure protein of SARS-Cov-2, the genome plasmid containing wildtype or truncated UTRs was added in equivalent molar ratios. Firefly luciferase were used as reporter of genome with wildtype UTRs, while renilla luciferase for truncated UTRs Subsequent to infection experiments with the packaged iVLPs, luminescence readings were acquired using the Duo-Lite Luciferase Assay System (Vazyme).

### Western blotting

An appropriate volume of supernatant containing iVLPs was combined with 1/4 volume of protein loading buffer and heated in a boiling water bath for 5 minutes. Post-cooling, the mixture was briefly centrifuged, and the resulting pellet was loaded onto a suitable tube. Electrophoresis was conducted on a gradient gel, followed by gel transfer to a PVDF membrane. The membrane was then blocked with a quick blocking solution and subsequently incubated with primary antibodies overnight at 4°C. After washing with TBST buffer, the membrane was incubated with secondary antibodies for 1 hour at room temperature. Chemiluminescence imaging was performed using iBright1500 (Invitrogen, USA).

### Digital Western blotting

Capillary-based Western blot (Wes) analysis was performed according to the manufacturer’s instructions for the 12-230 kDa protein separation module (#SM-W004; ProteinSimple). The supernatant containing iVLPs was centrifuged at 12,000 x g for 3 minutes at 4°C, and the supernatant was collected for use. The sample was diluted 1:4 with 10X sample buffer and PBS, mixed thoroughly, heated to 95°C for 5 minutes, and cooling on ice for 5 minutes to facilitate viral lysis. Sample preparation was carried out according to the manufacturer’s protocol, using a 5x mastermix without DTT, heated to 95°C for 5 minutes, and cooled on ice for 5 minutes. The prepared samples were loaded onto Simple Western plates. The primary antibody to SARS Spike Protein (1:1500, NB100-56578, Novus), and SARS-CoV-2 Nucleocapsid (1:80, 40588-T62, Sino Biological) was diluted in ProteinSimple’s Antibody Diluent 2. The secondary antibody and chemiluminescent substrate were used Anti-Rabbit Detection module (#DM-001; ProteinSimple). Proteins were quantified and visualized by “Compass for SW” software (ProteinSimple)

### Statistical Analysis

Statistical analysis was conducted using GraphPad Prism 9.5 software, employing an independent t-test to compare between two groups. A significance level of P < 0.05 was considered indicative of a statistically significant difference.

## Acknowledgements

We thank Dr. Yilong Yang, Dr. Xiaodong Zai, Dr. Yaohui Li for helpful discussion. We thank Yuemei Gao, Shuo Wang, Yihao Wei for their support.

## Authors’ contributions

KY, ZZ, FS, PL, Yue Z performed the experiments. ZZ, WS and KY performed statistical analyses. YZ,GY,JZ, JX and ZZ participated in the design of the study and wrote the manuscript. All authors read and approved the final version of the manuscript.

## Notes

### Competing Interest Statement

The authors have declared no competing interest.

