## Supplementary Figures for "Involvement of 5’ and 3’ UTRs on SARS-CoV-2 Genome Packaging"

**Supplementary figure. 1**

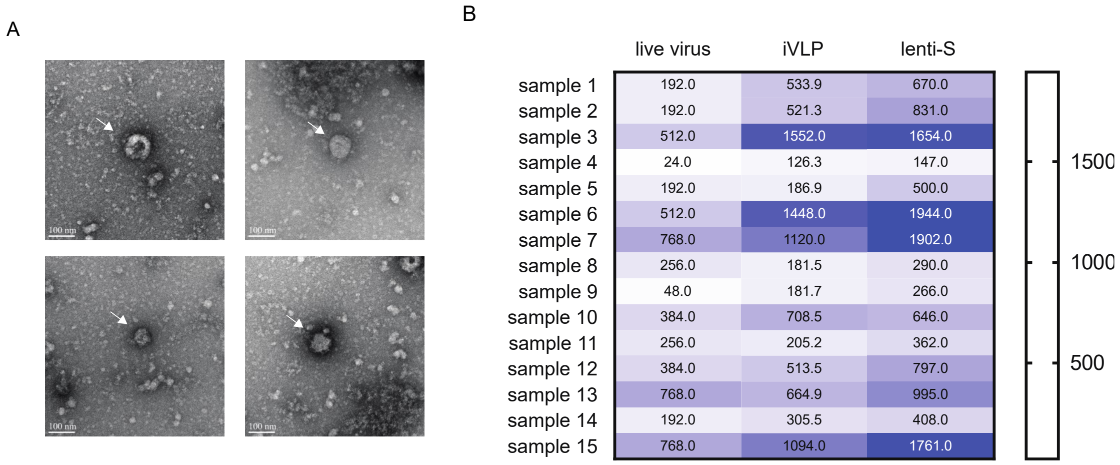


(A) Transmission electron microscope picture of iVLPs.

(B) Comparation of neutralization assay of vaccinated serum using live SARS-Cov-2, iVLPs design and generated in present, pseudotype lentiviral production decorated with spike protein of the viruses.

**Supplementary figure. 2**

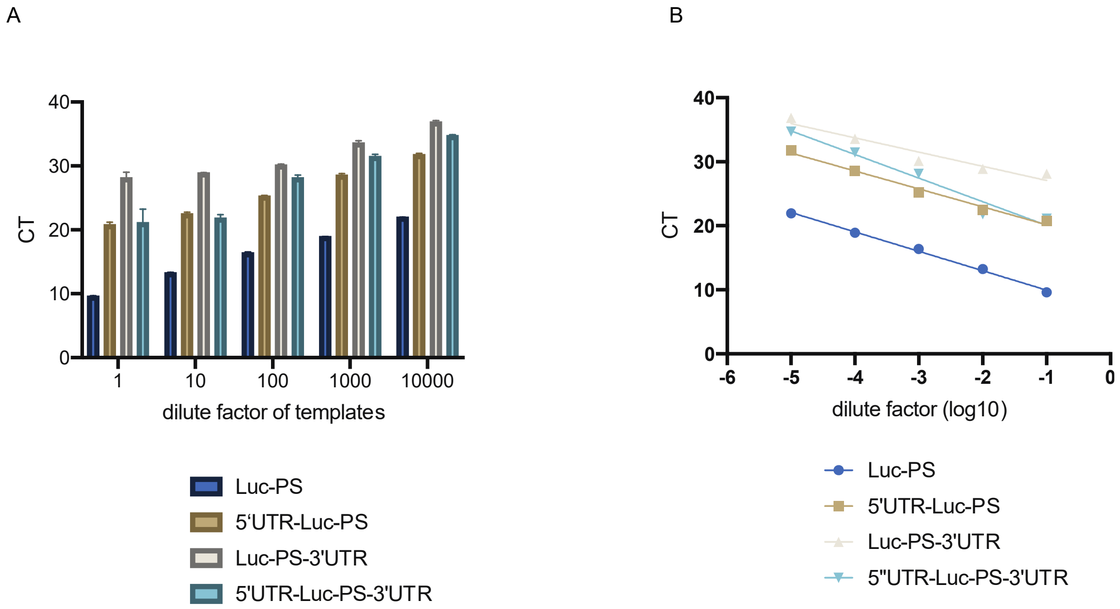


(A) qPCR detection of 4 different templates using primer set designed for specifically identify Luc-PS genome.

(B) qPCR detection of 4 serial diluted templates using corresponding primers sets.

**Supplementary figure. 3**

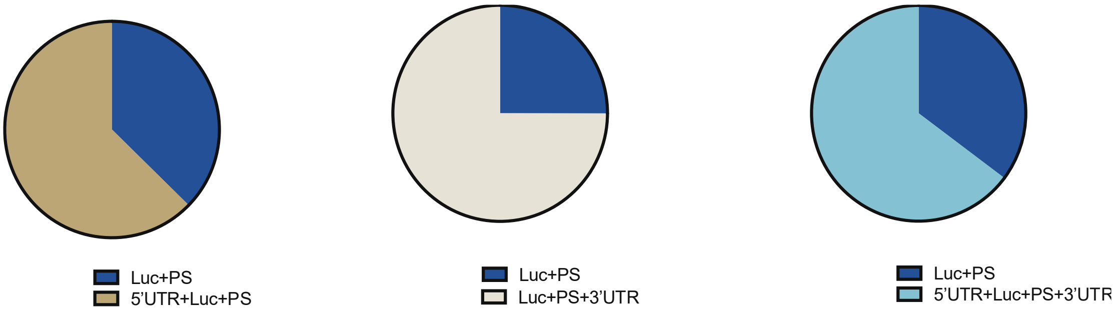


Relative portions of iVLP genomes with or without UTRs in pairwise competition assay.

**Supplementary figure. 4**

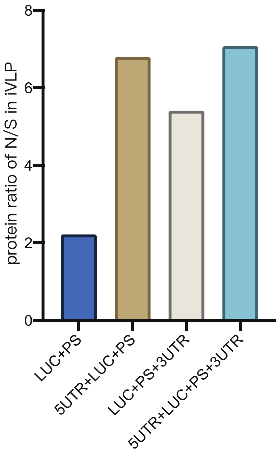


Relative expression of N protein in different iVLPs.

**Supplementary figure. 5**

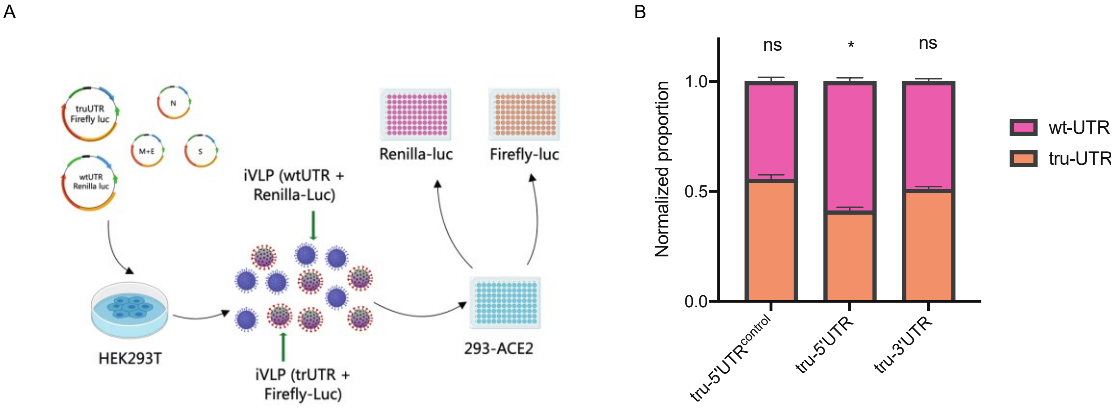


(A) Overview of competition test of iVLP genomes with wildtype or truncated UTRs.

(B) Relative portions of iVLP genomes with wildtype (wt) or truncated (tru) UTRs in competition assay.

**Supplementary table**

(1) EC50 of different iVLPs.

(2) Primers and probes used in the study. Underlines indicated homologous arms. Fwd, forward. Rev, reverse. Prb, probe.

(3) Linar regression of threshold cycle versus the dilution factor of template.

Supplementary table 1

| iVLPs | EC50 |
|---|---|
| Luc-PS | 414.3 |
| 5UTR-Luc-PS | 248.8 |
| Luc-PS-3UTR | 102.4 |
| 5UTR-Luc-PS-3UTR | 292.2 |

Supplementary table 2

| Primers | Sequence |
|---|---|
| Luc-PS-3UTR fwd | GCTGGCTAGCTTGTTAGACGAAGCTTGCCGCCATGGAAGATGCC |
| Luc-PS-3UTR rev | CTCCTTCTTAAAGGAGGCGGCC |
| tru-5UTR-1 fwd | CTGGCTAGCTTGTTAGACGAAGCTTCTTGTAGATCTGTTCTCTA |
| tru-5UTR-2 fwd | CTGGCTAGCTTGTTAGACGAAGCTTCATGCTTAGTGCACTCAC |
| tru-5UTR rev | CTGGCTAGCTTGTTAGACGAAGCTTCATGCTTAGTGCACTCAC |
| 5UTR fwd | CGGGTGTGACCGAAAGGTAA |
| 5UTR rev | GGGCCCTTCTTAATGTTTTTGG |
| 5UTR prb | ACCGGTCGCCGCCATGGA |
| 3UTR fwd | GGACTTGAAAGAGCCACCACAT |
| 3UTR rev | CATTGTTCACTGTACACTCGATCGT |
| 3UTR prb | TTCACCGAGGCCACGCGGA |
| Luc fwd | CTGGCTAGCGCCGCC |
| Luc rev | CCGTCTTCGAGTGGGTAGAATG |
| Luc prb | ATGCCAAAAACATTAAGAAGGGCCCAGC |

Supplementary table 3

| | Luc-PS | 5'UTR-Luc-PS | Luc-PS-3'UTR | 5"UTR-Luc-PS-3'UTR |
|---|---|---|---|---|
| Equation | Y = -3.030*X + 6.926 | Y = -2.800*X + 17.36 | Y = -2.221*X + 24.85 | Y = -3.685*X + 16.39 |
